# Functional arrays of human pluripotent stem cell-derived cardiac microtissues

**DOI:** 10.1101/566059

**Authors:** Nimalan Thavandiran, Christopher Hale, Patrick Blit, Mark L. Sandberg, Michele E. McElvain, Mark Gagliardi, Bo Sun, Alec Witty, George Graham, May Mcintosh, Mohsen A. Bakooshli, Hon Le, Joel Ostblom, Samuel McEwen, Erik Chau, Andrew Prowse, Ian Fernandes, Penney M. Gilbert, Gordon Keller, Philip Tagari, Han Xu, Milica Radisic, Peter W. Zandstra, Dana Nojima, Hugo Vargas, Yusheng Qu, Alykhan Motani, Jeff Reagan

## Abstract

To accelerate the cardiac drug discovery pipeline, we set out to develop a platform that would be amenable to standard multiwell-plate manipulations and be capable of quantifying tissue-level functions such as contractile force. We report a 96-well-based array of 3D human pluripotent stem cell (hPSC)-derived cardiac microtissues - termed Cardiac MicroRings (CaMiRi) - in custom printed multiwell plates capable of contractile force measurement. Within each well, two elastomeric microcantilevers are situated above a ramp. The wells are seeded with cell-laden collagen which, in response to the slope of the ramp, self-organizes around tip-gated microcantilevers to form contracting CaMiRi. The contractile force exerted by the CaMiRi is measured and calculated using the deflection of the cantilevers. Platform responses were robust and comparable across wells and we used it to determine an optimal tissue formulation. We validated contractile force response of CaMiRi using selected cardiotropic compounds with known effects. Additionally, we developed automated protocols for CaMiRi seeding, image acquisition, and analysis to enable measurement of contractile force with increased throughput. The unique tissue fabrication properties of the platform, and the consequent effects on tissue function, were demonstrated upon adding hPSC-derived epicardial cells to the system. This platform represents an open-source contractile force screening system useful for drug screening and tissue engineering applications.

## INTRODUCTION

One significant bottleneck in engineered heart tissue screening is the high cost associated with the large cell numbers required for high-fidelity in vitro cardiac models. Miniaturization of self-organizing cardiac organoid generation is one approach to address this bottleneck which has been previously applied to tissue engineering using photolithography- and micromachining-based fabrication techniques ^1–9^. Unfortunately, these systems are rarely easy to manufacture on a large scale and production often requires a high level of skill. Furthermore, these methods are time consuming in the fabrication and seeding stages of tissue building, restrictive to certain design geometries (for instance vertical walls spanning either the micrometer or centimeter scale but not both), and expensive to prototype. Another challenge typically associated with these microfluidic- and microstructure-based devices is adaptation to existing tissue culture equipment such as microscope stage adapters and liquid dispensing tools. Potential complications may also arise upon the introduction of cells into these devices, including a tissue transfer step to another device after tissue formation for long term culture ^8, 10–13^, or a transfer step to a measurement device such as a force transducer ^7, 9, 14, 15^ to measure contractile force. These steps not only complicate and extend the overall process but may also introduce unwanted variability in baseline tissue conditions, a major hurdle in designing quality screening applications. Additionally, contractile force, a key functional metric in heart health, is not typically measured in engineered heart tissue screening platforms, limiting the functional assessment and predictive ability of these platforms ^1, 2^.

We previously determined key design criteria for the formulation of human pluripotent stem cell (hPSC)-derived cardiac microtissues including an optimal ratio of input cells for tissue formation and induction of intratissue cell alignment ^3^. Here we outline the incorporation of these criteria in the design of a 3D printing based 96-well plate screening platform that supports both cardiac microtissue self-organization and contractile force measurement. Our new platform is simple to manufacture and is seeded with cells that form Cardiac MicroRings (CaMiRi) - cardiac microtissue that is configured for contractile force measurements, in addition to the standard calcium handling and conductance parameters conventionally measured in engineered cardiac tissue ^3^. Notably, to maximize accuracy and reproducibility, we have designed a simple process for measuring contractile force *in situ* with automated data collection using only a bright field camera. Briefly, CaMiRi are formed by seeding cell-laden collagen into a reservoir in each well. The cell-laden collagen forms an organoid around two elastomeric microcantilevers located at the base of each well. The contractions of the CaMiRi cause the microcantilevers to deflect towards the center of the well. The displacement of this deflection can be imaged via a digital camera in each well and used to calculate a total magnitude contractile force, using both the structural and material properties of the elastomer microcantilevers. We optimized this platform by conducting a Central Composite Design (CCD) experiment to test the effects of several input variables on the contractile force exerted by the CaMiRi. We then validated our platform with a cardiotoxicity study using compounds with known effects on cardiomyocyte contractile force response and automated key bottlenecks in the experimental process to enable high throughput measurement of contractile force. Finally, we added fractions of hPSC-derived epicardial cells to our optimized cardiac tissue formulation to test effects on contractile force in CaMiRi. This platform provides a widely accessible solution for functional screening and validation of cardiac-associated drugs.

## RESULTS

### Development of 3D printing-based method to produce 96-well plates for Cardiac MicroRing (CaMiRi) formation

We designed a platform which would allow us to culture and measure the contractile forces exerted by cardiac microtissues. The platform consists of wells containing dual polydimethylsiloxane (PDMS) cantilevers around which CaMiRi form due to the compaction of a collagen 1-based matrix by the cardiomyocytes and cardiac fibroblasts (cFB) contained within (Figure 1A). Our elastomeric microcantilevers provide resistance during auxotonic contraction ^16^ and serve as sensors for measuring contractile force ^1, 6, 17^. To build this construct, we designed and rendered a 3D Computer Aided Design (3D CAD) of the 96-well tissue culture plate model with outer dimensions and inter-well distances based on Costar™ 96 well plates to be compatible with conventional multi-well cell culture and imaging equipment, such as multi-well pipettors and automated imaging microscope systems (Figure 1A and B). This design was printed using an Objet 30 Pro 3D printer to generate a positive mold. Next, we produced a negative PDMS mold which was used to produce a complementary positive PDMS mold. The positive PDMS mold was then used to create a polyurethane-based negative master mold which can generate multiple PDMS plates. This process can be used to create 96-well plates with any desired geometry. The open-source designs for our plates are available in the Supplementary Materials for researchers to further explore and build customized designs. The final device and micro-cantilevers (Figure 1C and D) were tested with a force transducer (Microsquisher) to determine a force-displacement standard curve for force measurements. The theoretical curve (Supplementary Figure 1) for this relationship was determined and confirmed empirically (Figure 1E).

**Figure 1.**
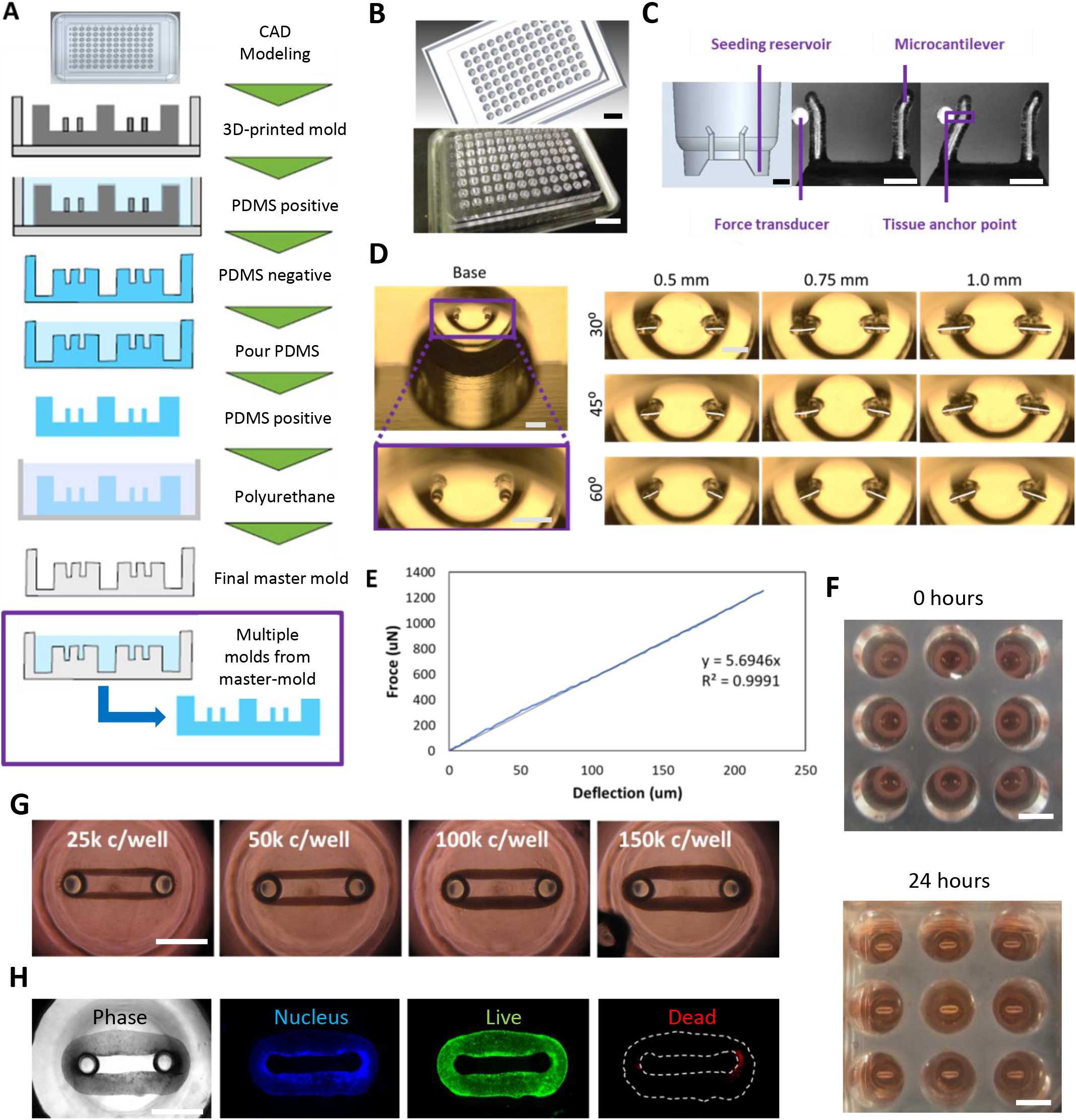
Design and manufacture of Cardiac MicroRing (CaMiRi) platform. (A) Process flow of substrate manufacturing. The process started with creating a three-dimensional Computer Aided Design (3D CAD) model of the plate. The design is then printed using an Objet 30 Pro 3D printer to generate a positive mold. A negative PDMS mold is then created which is followed by a complementary positive PDMS mold. This positive PDMS mold is used to then create a polyurethane-based negative master mold which can be used to mold multiple PDMS plates. (B) 3D CAD model of a plate for 3D printing (top panel) and actual 3D-printed mold (bottom panel). Scale bars represent 20 mm. (C) Cross section of a well showing the seeding reservoir where cells are pipetted and the microcantilevers used for force-displacement measurement (empirically calibrated with a force transducer (Micro-Squisher) with tissue anchor points to hold microtissue in place. Scale bars represent 1 mm. (D) Tissue anchor angles and lengths were tested to determine optimal anchor geometry. Scale bars represent 1 mm. (E) Force-displacement curve of cantilevers as measured with the Micro-Squisher. (F) Cell-laden collagen seeded into reservoirs of CaMiRi device remodel into cardiac tissues tethered around microcantilevers within 24 hours. 9 wells of a 96-well plate containing CAMIRI are also shown. Scale bars represent 5 mm. (G) CaMiRi formulated from a range of total input cells (25,000; 50,000; 100,000; and 150,000 cells seeded per well. Scale bars represent 1 mm. (H) Live/dead staining of CaMiRi. From left to right: bright field, nuclei in blue, live cells in green, and dead cells in red. Scale bars represent 1 mm.

We designed the well reservoir volumes to precisely match the cell seeding volumes to facilitate the formation of a fully closed ring in the reservoir which is critical for proper tissue formation (Supplementary Figures 1 and 2). To ensure that any residual debris from the cell seeding step would not interfere with the formation and maturation of the CaMiRi, and to maximize tissue surface area exposed to culture medium (which is important for nutrient diffusion), we designed our wells to promote passive movement of the self-organized CaMiRi away and upward from the point of initial tissue formation and into a toroidal geometry (Supplementary Figure 3). To this end, we added a ramp in the middle of the cell seeding reservoir to promote passive tissue movement away from the reservoir during the tissue compaction phase and toward the vertical tip of the cantilevers. To prevent the CaMiRi from slipping off the cantilevers and to ensure that the fully formed CaMiRi would be positioned at the same vertical location on the cantilevers during contractile force measurement, we incorporated an anchor by angling the cantilever tips away from the well center (Figure 1C). The consistent positioning of the CaMiRi is critical as the absolute magnitude force measured via cantilever deflection is a function of the vertical positioning of the contracting CaMiRi. The anchor length and angle selected had to be sufficient to trap the tissue yet not too long or acute so as to not survive the demolding process. We screened three angles (30 degrees, 60 degrees, and 90 degrees to the horizontal) and 3 anchor lengths (0.5 mm, 0.75 mm, and 1.0 mm) (Figure 1D) in four demolding tests and found only the 0.5 mm anchors with either 60 or 45 degree angles remained fully intact. We chose the 0.5 mm length anchors with the 45 degree (more acute) angle to achieve better tissue anchorage.

**Figure 2.**
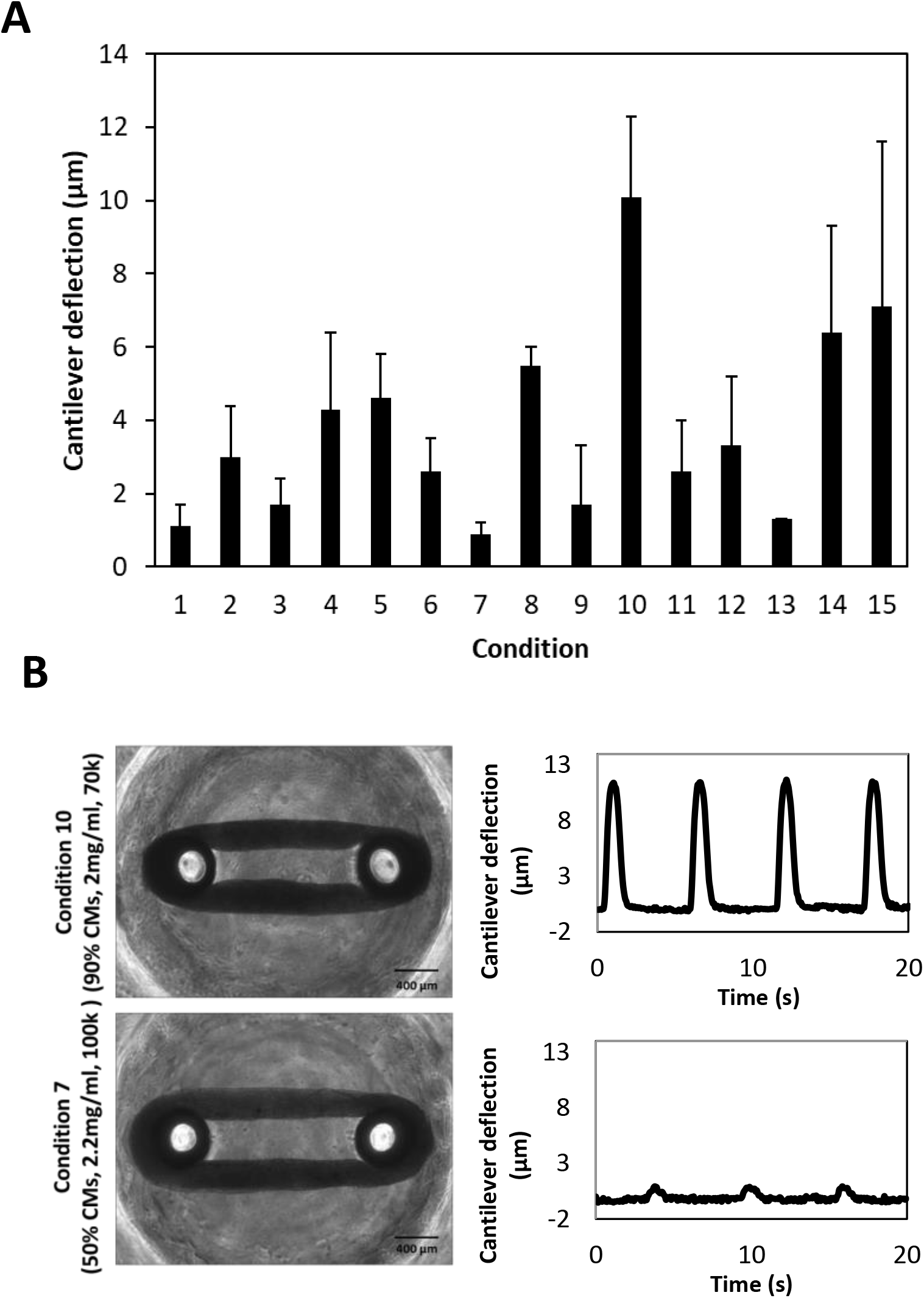
Central Composite Design (CCD)-informed formulation of CaMiRi. (A) Contractile force (as measured by cantilever deflection) of cardiac microtissues in all tested conditions, with condition 10 resulting in the highest measured contractile force. Data are reported from three independent experiments, each with three replicates, as the mean ± standard deviation. (B) Representative plots of cantilever deflection of optimal and least optimal conditions. Tissue images are shown in bright field. Scale bars represent 400 µm.

### Formation of CaMiRi in 96 well plate platforms suitable for contractile force measurements

Next, we developed a seeding protocol for the formation of CaMiRi amenable to high-throughput robotic pipetting systems. First, chilled cell-laden collagen 1-based extracellular matrix is pipetted into the wells of the plate (Supplementary Figure 3). Within 24 hours, the collagen polymerizes and a CaMiRi forms around the two elastic micro-cantilevers located at the base of each well (Figure 1F). As the CaMiRi contracts around the micro-cantilevers, the contractile force deflects the cantilevers. The displacement of the deflection can be video imaged in each well and custom edge-detection software can be used to analyze and calculate a total contractile force, using both the structural and material properties of the micro-cantilevers (Supplementary Figure 5).

We tested the effect of varying input cell numbers on CaMiRi formation to test the capability of forming a range of tissue sizes. Cell input numbers ranging from 25,000 to over 150,000 cells per tissue were tested (Figure 1G). The cells in the CaMiRi maintained viability over the course of remodeling (Figure 1H).

To maximize successful CaMiRi formation and tissue force generation, we next optimized our seeding protocol with respect to cardiomyocyte composition, collagen concentration and seeding density. These factors were assessed using a three-level face centred central composite design (CCD) which allowed us to create a linear regression model to obtain a range of response values for each tested factor. The range of values selected in the CCD for these three factors were based upon previous studies in the formulation and function of cardiac microtissues ^1, 3, 6, 18^. Specifically, the CCD explored input cardiomyocyte compositions of 50%, 70%, and 90% (with the remaining tissue composition consisting of cFB acquired from Promocell); collagen concentrations of 1.8 mg/mL, 2.0 mg/mL, and 2.2 mg/mL; and total cells per CaMiRi of 40,000; 70,000; and 100,000 (Supplementary Figure 6A and B). A quadratic model was selected to fit the data and an ANOVA was performed to assess the significance of each term as a means of maximizing successful CaMiRi formation and force generation.

The fifteen seeding conditions exhibited similar contraction frequencies (Supplementary Figure 6C) but varied in magnitude of micro-cantilever deflection (Figure 2A and B). Condition 10 (90% cardiomyocytes, 2 mg/mL collagen and 70,000 total cells) exhibited the highest mean micro-cantilever deflection with a value of 10.1 ± 2.2 µm, while condition 7 (50% cardiomyocytes, 2.2 mg/mL collagen and 100,000 total cells) resulted in the lowest mean micro-cantilever deflection at 0.9 ± 0.3 µm. Overall, the CaMiRi with the highest composition of cardiomyocytes (90%) and medium levels for collagen concentration (2.0 mg/mL) and total cells per CaMiRi (70,000) exhibited the highest contractile force.

Contraction frequency data were used to generate response surface models to determine the optimal formulation that would lead to maximal contractile force and dynamic measurement range (Supplementary Figure 7). Fitting a quadratic model to the data demonstrated that the percentage of cardiomyocytes seeded (p < 0.05) and the square of collagen concentration (p < 0.01) were statistically significant. From this model, the condition leading to maximum micro-cantilever deflection was predicted to be 90% cardiomyocytes, 2.03 mg/mL collagen, and 75,400 input cells per well. For practicality, a collagen concentration of 2.0 mg/mL and a cell concentration of 75,000 per well were subsequently used. Whole mount immunostaining of least optimal condition (7), optimal condition (10), and predicted optimal condition CaMiRi was performed to observe cardiac troponin T alignment, which was most strongly displayed in the optimal CaMiRi condition (Figure 3).

**Figure 3.**
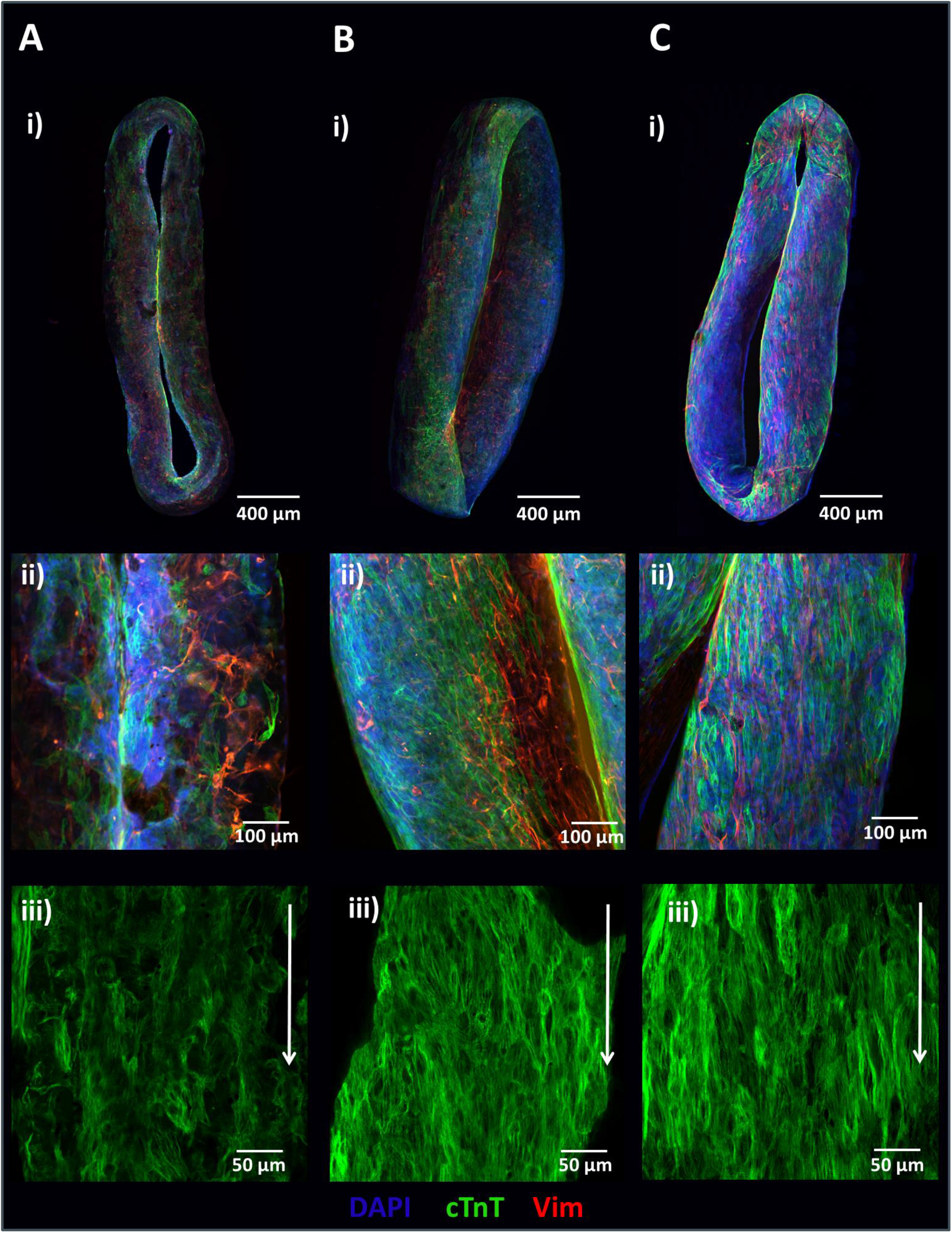
Tissue structure of CaMiRi. (A) Whole mount CaMiRi immunostaining of least optimal condition 7, (B) condition 10, and (C) the optimal condition. DAPI shown in blue, cardiac troponin T (cTnT) shown in green, and vimentin shown in red. Middle row (ii) shows higher magnification of CaMiRi. Final row (iii) shows cTnT in green aligning in parallel with white arrows. Scale bars represent 400 µm, 100 µm, and 50 µm, of images in top, middle, and bottom rows, respectively.

### Assessing the CaMiRi platform to measure contractile force response to cardiotropic drugs with known effects

To test the consistency of our platform for measuring contractile force, we studied CaMiRi response to cardiotropic drugs. We first generated CaMiRi based using the optimized seeding conditions determined from our CCD experiment (Supplementary Figure 3A). The tissues were cultured for 2 weeks before drug screening assays were conducted. We chose 3 cardiotropic drugs with known effects: Blebbistatin and Nifedipine are negative inotropes which decrease contractile force, and Isoproterenol is a positive inotrope which increases contractile force. For each drug, we tested 5 doses based on previous studies using these drugs ^19, 20^. The three drugs were administered and tissues were incubated for 20 minutes and then assayed relative to a vehicle control containing 0.1% DMSO.

Blebbistatin, a potent inhibitor of myosin, has been previously shown to substantially reduce the force of contraction of explanted rabbit myocardial tissues, with little to no effect on heart pacing ^21^. Increasing concentrations of Blebbistatin resulted in diminishing contractile force responses in the CaMiRi (Figure 4A). Several studies have demonstrated that the administration of Isoproterenol, an adrenoreceptor agonist, causes an increase in the force of contraction of myocardial tissues ex-vivo ^22, 23^. In contrast to Blebbistatin, administration of Isoproterenol led to a gradual increase in observed contractile force, which peaked at 1 µM (Figure 4B). Finally, Nifedipine, an L-type Ca2+ channel blocker, caused a decrease in relative contractile force (Figure 4C) with increasing dose, consistent with results reported in ex vivo myocardial tissues ^24^ and clinical trials ^25^. Overall, our results are consistent with previously reported electrophysiology analysis using the xCELLigence and microelectrode arrays (MEAs) systems on iCell cardiomyocytes ^20^. We also used the CaMiRi system to measure the effect of 7 growth factors with reported effects on cardiomyocytes on two electrophysiological parameters of cardiac tissue: maximum capture rate (a measure of tissue response to stimulation) and excitation threshold (a measure of tissue excitability). Although there were no significant effects relative to control on maximum capture rate, we observed that both Insulin Growth Factor (IGF)-1 and Histidine-glyco-protein (HRG) elicited significant decreases in excitation threshold relative to control (p < 0.05). These growth factors could thus be the basis for a drug to treat conduction block in patients with problematic electrophysiology. There was no noticeable change in either response to the remaining 5 growth factors (Supplementary Figure 8A and B).

**Figure 4.**
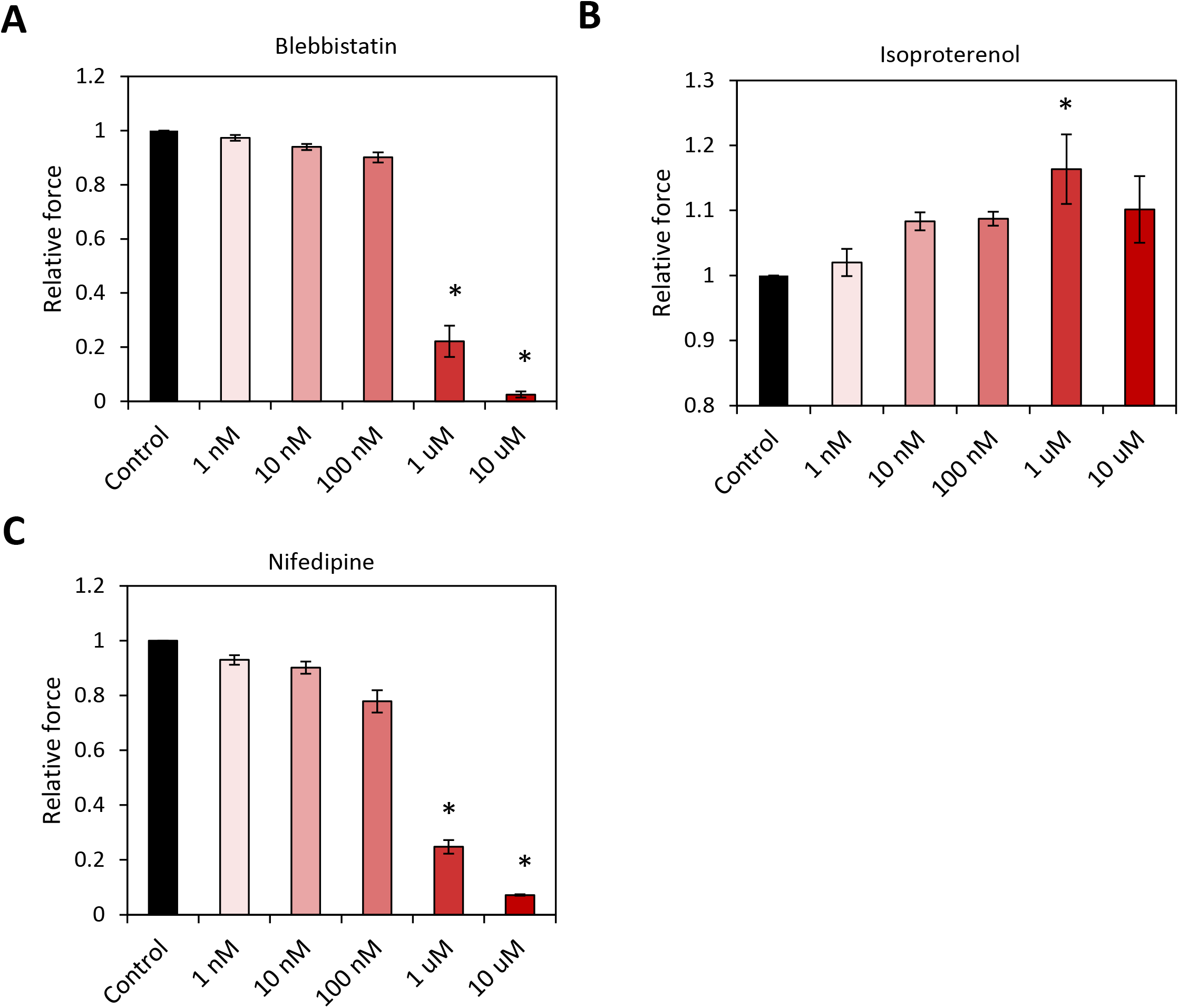
CaMiRi response to cardiotropic drugs and growth factors. Drugs were administered and tissues incubated for 20 minutes before contractile force assay. Along with the vehicle control, five drug concentrations of 1 nm, 10 nm, 100 nm, 1 µm, and 10 µm are administered. Data shown are relative change in contraction amplitude due to the administration of (A) Blebbistatin (negative inotrope), (B) Isoproterenol (positive inotrope), and (C) Nifedipine (negative inotrope), as compared to the vehicle control. Data are reported from three independent experiments, each with three replicates, as the mean ± standard deviation. *P < 0.05.

### Assay automation for high throughput measurement of contractile force

To address experimental bottlenecks, we developed automation protocols to accelerate and standardize CaMiRi seeding, image acquisition, and image analysis. Cell-laden collagen was dispensed in discrete droplets by an Agilent Bravo robotic liquid handler from a source plate to a destination plate (Supplementary Figure 9A and B), both maintained at 4°C to avoid gelation. Droplets were then driven to well bottoms via cooled centrifugation, again at 4°C (Supplementary Figure 9C). Centrifugation was optimized at 200 g for 5 minutes, both qualitatively by visual inspection of ring formation and functionally by maximizing post deflection (Supplementary Figure 9D). Collagen concentration was also increased to 2.2 mg/mL to promote uniform CaMiRi generation. Uniform CaMiRi subsequently developed in 96 well format within 7 days (Supplementary Figure 9E).

Automated imaging of CaMiRi in the 96 well PDMS plate presented significant challenges due slight spacing between wells and minor variability in absolute post heights relative to the plate bottom, both due to the variation involved in plate curation. To overcome these issues, we employed a targeted imaging approach on the microscope in which wells were first imaged at low magnification to identify post coordinates. An image-based autofocus routine was then performed at high magnification, and videos of post edges were acquired at a high frame rate which were used to produce kymographs (Figure 5A). This system, equipped with environmental control and liquid handling capabilities, also enabled automation of compound addition studies, reducing experiment workflow from five to one hour of hands-on time. Automated image analysis was also employed to accelerate analysis of post deflection videos.

**Figure 5.**
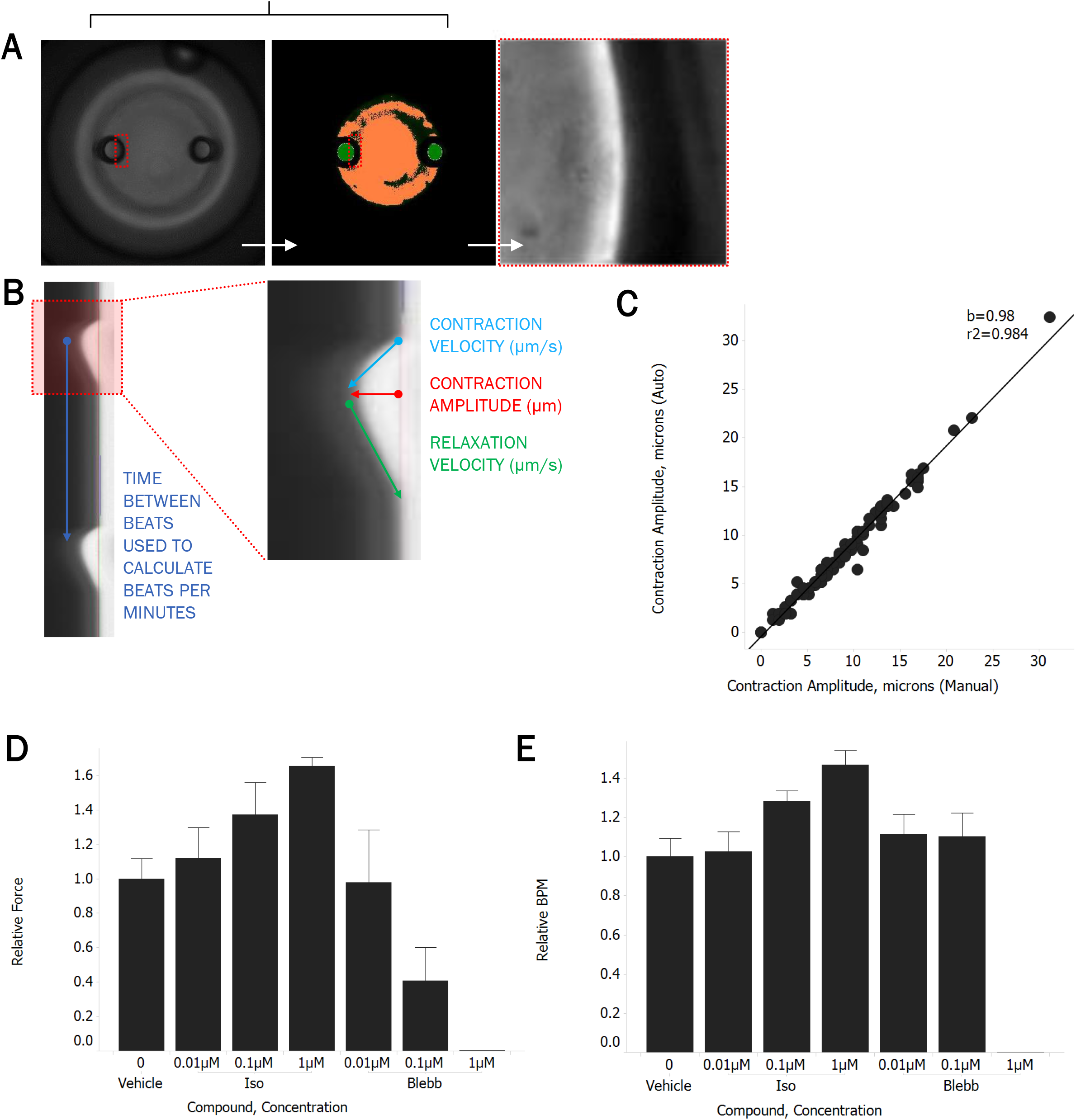
Automated CaMiRi imaging and analysis. (A) Whole well imaged with 2X objective on Molecular Devices Image Xpress Micro 4 under transmitted light (left), after image analysis routine to identify posts (center), and at 20X after identification of right edge of left post (right). (B) Kymograph of post deflection over time, noting calculation of output parameters. (C) Correlation between contraction amplitudes calculated with automated script (y-axis) vs. manual calculations (x-axis) performed on kymographs. (D, E) Response of auto-seeded CaMiRi to isoproterenol (green) and blebbistatin (red). Data are reported from a single trial as the median ± median absolute deviation.

Kymographs were generated from video streams and analyzed in a custom MATLAB (Mathworks) program to export beats per minute, contraction amplitude, contraction velocity, and relaxation velocity for each CaMiRi (Figure 5B). Image analysis software was designed to achieve a high degree of correlation between automated and manual measurements over a wide range of post deflection amplitudes (Figure 5C). Finally, we validated our automated workflow by confirming positive and negative inotropic responses of isoproterenol and blebbistatin, respectively (Figure 5D), which were in line with the expected range of effect, along with relative beating frequency (Figure 5E).

### Using the CaMiRi system to rapidly explore cell mixing studies on tissue formation

We next sought to engineer a CaMiRi composed of hPSC-derived epicardial and hPSC-CM layers to represent a more anatomically-relevant cardiac tissue ^26^. In the mammalian heart, the epicardium is located on the outer region of the heart, facing inward to the pericardium and the pericardial cavity (Figure 6A). Epicardial cells are present in both the adult, where they give rise to cell types such as the cFB that are critical during both homeostasis and repair, and the developing heart ^26^. Epicardial cells secrete factors that promote cardiomyocyte hyperplasia and hypertrophy in the myocardium, which in turn leads to an increase in contractile force. To determine the effects of epicardium on cardiomyocyte contractility in our CaMiRi, we received hPSC-derived epicardial cells using a published 31-day differentiation protocol ^26^. The cells produced in this protocol display epicardial-like morphological characteristics and express markers of the epicardial lineage (WT1, TBX18, and the retinoic acid– producing enzyme ALDH1A2 (aldehyde dehydrogenase)) ^26^.

**Figure 6.**
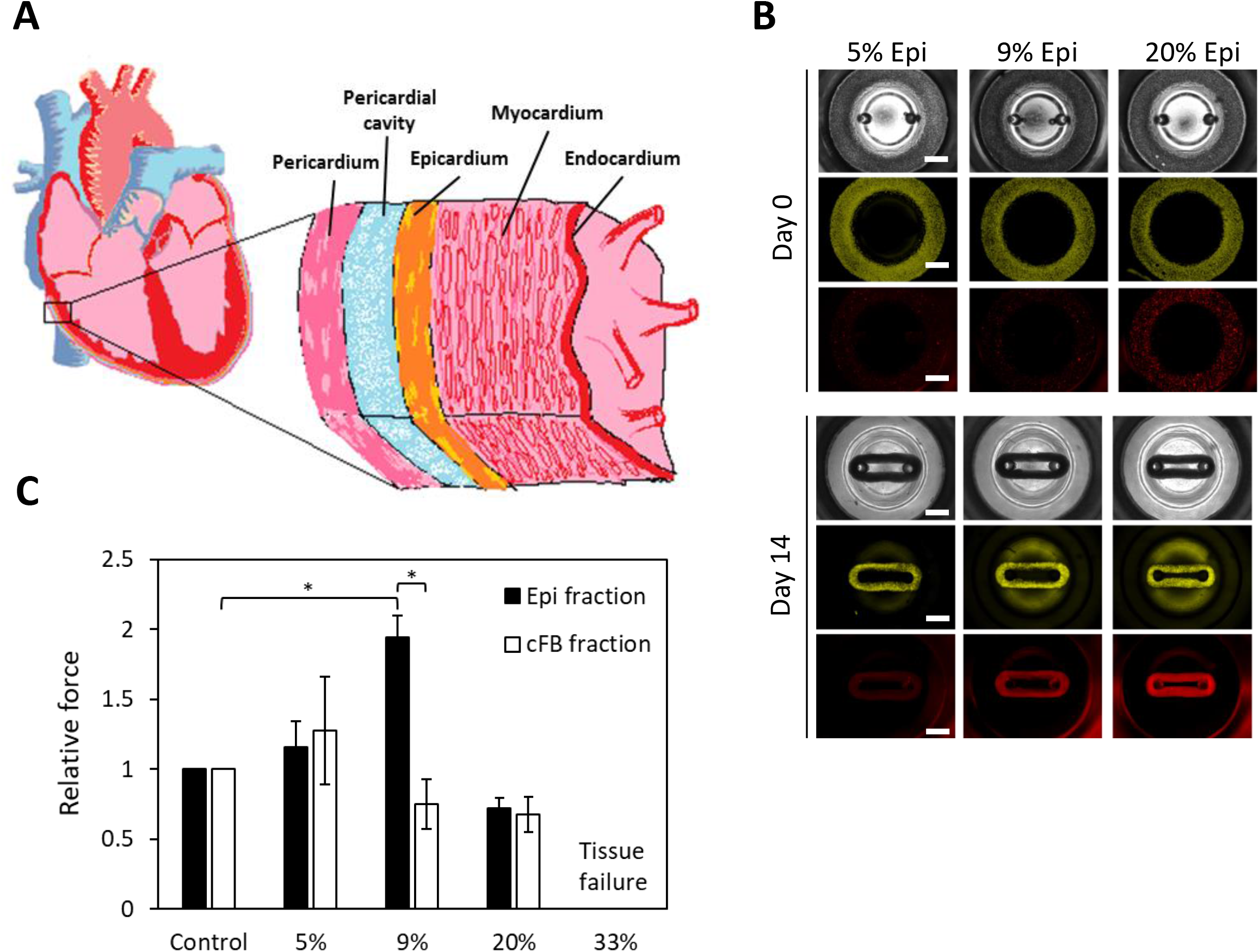
Exploring the effects of epicardial cells in CaMiRi. (A) Cross-sectional schematic of heart wall including epicardium, myocardium, and endocardium shows the complexity of cell types that compose heart wall tissue. (B) Spatial organization and tissue compaction of hPSC-CM and epicardial cells in CaMiRi (top panel is day 0, and bottom panel is day 14). Epicardial fractions of 0% (control), 5%, 9%, 20%, and 33% were used to create tissues. In grayscale (top layer of each panel), bright field is shown of tissues. Green (middle layer) represents hPSC-CM and red (bottom layer) represents hPSC-derived epicardial cells. Scale bars represent 1 mm. (C) Contractile force readings of CaMiRi composed of five different epicardial cell fractions along with associated cFB controls. CaMiRi composed of 33% epicardial fraction failed due to overwhelming tension forces. Data are reported from four independent experiments, each with two to four replicates, as the mean ± standard deviation. *P < 0.05.

We first labeled hPSC-CM with cell tracker green CMFDA and epicardial cells with cell tracker red CMTPX. We then dosed in epicardial cell fractions (5%, 9%, 20%, and 33%) into our optimized base CaMiRi formulation to determine its functional effects. We also conducted a control set in parallel where additional cFB fractions (5%, 9%, 20%, and 33%) were integrated into the optimized base CaMiRi formulation. The mixed cell formulations were seeded (Figure 6B upper panel) and after 2 weeks (Figure 6B bottom panel) CaMiRi formation was observed. We measured contractile force for each condition and observed maximum contractile force in CaMiRi composed of 9% epicardial cell fraction (Figure 6C). The force generated in CaMiRi composed of 9% epicardial cell fraction was ∼2-fold higher than the associated controls. Epicardial cells do not contribute directly to the active contractile force in cardiac tissue ^26^, suggesting their interaction with hPSC-CM leads to larger force output of the hPSC-CM or they affect the transmission of force via structural changes in the tissue. As the fraction of both epicardial cells and cFB increased, specifically in the 20% and 33% fractions, contractile force lowered, and instances of tissue failure increased. In the 33% condition, all tissues failed due to tension-induced rips. Optimization of cardiac tissue formulation using epicardial cells and eventually including other stromal cell types will be critical in creating physiologically relevant and highly predictive models. Our platform has been shown to facilitate these studies effectively.

As a final step, we attempted to create a process to build a more complex tissue with the goal of understanding the limits of creating sophisticated models to explore complex biology in our platform. We did this by taking advantage of the well reservoir that becomes vacated by the tissue as it forms and moves along the micro-cantilever to embed within the primary CaMiRi with a secondary cell type (Supplementary Figure 10). As a proof-of-principle experiment, we first seeded our platform with Green Fluorescent Protein (GFP)-expressing fibroblasts, which formed a ring around the dual micro-cantilevers, and subsequently seeded Red Fluorescent Protein (RFP)-expressing fibroblasts into the reservoir. The RFP+ cells formed a secondary tissue layer, also in ring form, around the initial GFP+ ring to create a tissue with distinct cell type separation with minimal cell type mixing (Supplementary Figure 10). This sequential seeding method not only demonstrates that our platform can be used to create more complex tissue structures compared to simultaneous seeding methods, but also provides a model to study tissue-tissue interactions.

## DISCUSSION

In this work, we developed an engineered cardiac microtissue platform that now enables high-throughput contractile force measurements in a 96-well plate configuration. Our CCD experiment identified that the optimal conditions to form CaMiRi involved seeding 75,000 cells composed of 90% CM and 10% cFB in a matrix of 2.0 mg/mL of Collagen 1. It is important to note that the CM:cFB ratio identified here varies from our previous studies ^3^. This discrepancy is perhaps due to the nuanced functional differences of the input cell types resulting from the differentiation protocol used to generate the hPSC-CM or primary tissue the input cells are derived from.

Interestingly, the adult mammalian heart is composed of 20-30% cardiomyocytes and 70-80% non-myocytes ^27^. However, in newborn hearts, the percentage of cardiomyocytes are much higher at nearly 70% ^28^. The hPSC-CM we use represent an embryonic stage of development, providing a possible explanation for the higher percentages of CM needed for optimal contractile function in our CaMiRi. Perhaps once there are protocols to efficiently mature hPSC-CM to approach the morphology and phenotype that more closely resembles mature heart cells, the optimal ratios in our engineered tissues will also more closely resemble ratios found in native heart tissue.

Although our CaMiRi exhibited the expected responses to the cardiotropic drugs tested, greater changes in contractile force may be detected if the microtissues we generated possessed a mature electrical pacing apparatus. Interestingly, in a study of five patients, Ross et al. demonstrated that the infusion of isoproterenol, a positive inotrope, causes an increase in heart rate but with inconsistent effects on stroke volume ^29^. However, when the heart rate was held constant through electrical stimulation, the infusion of isoproterenol successfully provoked an increase in stroke volume in all patients. Thus, the lack of an even larger increase in measured contractile force in our platform due to isoproterenol treatment may result from a lack of mature pacemaker cells or external electrical stimulation during our assay.

It is important to note that observed effects of drug exposure are also dependent on the assay technology employed and its readout. For example, Micro Electrode Arrays (MEA) have been used to assess changes in electrical field potentials of electrically coupled populations of cardiac cells as a surrogate measure for the changes in the force of contraction. Likewise, the xCELLigence system uses changes in impedance measurements associated with electrically coupled cardiac cells as a surrogate for changes in force of contraction. Using these two assay platforms to assess the effect of isoproterenol on hPSC-CM monolayers, Guo et al. did not observe an increase in calcium flux electrical activity with the MEA system but did observe an increase in impedance on the xCELLigence system ^20^. In contrast to both technologies, the CaMiRi platform enables a direct measurement of contractile force. Furthermore, we have developed an automation pipeline to enable high throughput measurement of contractile force in a 96-well format.

Our studies with the epicardial and myocardial mixed tissues show the ability of our system to study cell-cell interactions within a tissue setting. Although our tissues lack spatial organization between the myocardial and epicardial fractions, we showed the possibility of using sequential seeding to study such phenomenon in a sequential tissue layering CaMiRi model. We envision that this concept will help with tissue studies in many systems that involve 2 or more functional cell types that interact either through cell-cell contact, secreted factors, or even through cell migration via chemokines.

We are interested in further investigating the effect of the epicardial fraction on contractile force in our CaMiRi platform. The epicardium may be mediating this effect via three potential mechanisms: either by providing structural support for contractile force transmission within the tissue; through the secretion of soluble factors that directly affect the CM fraction within the microtissues; or by providing additional fibroblasts following epithelial–mesenchymal transition. With respect to the second potential mechanism, many factors have been shown to be secreted by epicardial cells including IGF ^26^. We observed the positive effect of IGF on cardiac electrophysiological function, specifically on the excitation threshold of cardiac microtissues which is a measure of tissue excitability ^30^, which may play a role in this case as well, to increase contractility.

In conclusion, we have validated the advantages and demonstrated the potential of our simple-to-use and rapid-to-manufacture platform for studying engineered heart tissue. We hope this new method sets a new standard for cardiac tissue engineering and contractility testing, along with further possibilities for engineered custom geometry tissue culture plates applied to other tissue types and applications.

## METHODS

### Fabrication of Cardiac MicroRing (CaMiRi) 96-well plates

A modified 96-well plate with embedded microcantilevers was designed using Solid Edge Three-Dimensional Computer Aided Design (3D CAD) software (Siemens, Student Edition) and printed with an Objet 30 Pro 3D printer (Stratasys). A polydimethylsiloxane (PDMS) negative was molded from the 3D printed plate, and then a positive PDMS plate molded following it. Next, SmoothCast310 polyurethane (Smooth-On) was used to fabricate a master mold. CaMiRi PDMS devices were fabricated by pouring Sylgard 184 (Dow Chemicals) into the polyurethane mold, followed by degassing to remove bubbles and curing for 48 hours at 37°C. The resulting PDMS CaMiRi device was then carefully removed from the mould and visually inspected for defects with a Leica MZ6 stereomicroscope.

### Cardiac differentiation of human Pluripotent Stem Cells

Cardiac differentiation of HES2 (ES Cell International) human embryonic stem cells (hESC) was carried out as previously reported ^3^. The hESC were maintained and expanded as described previously ^3^. Briefly, HES2 cells were cultured on mouse embryonic feeders (MEF) for 6 days in HES2 maintenance media (80% DMEM/F12, 20% KOSR, 20 ng/mL bFGF, 0.5% P/S, 1% NEAA, 1% BME) and media was changed daily. Cells are maintained in normoxia at 37°C in a 5.0% CO_2_ atmosphere. The cells were then trypsinized along with MEFs and plated onto Matrigel (diluted at 1:30) coated plates at a split ratio of 1:3 for MEF depletion. After two days of MEF depletion, HES2 cells were again trypsinized and seeded into AggreWells manufactured in-house to form hEB. The hEB were generated using 400 µm microwell PDMS inserts cast from a silicon master mould. The inserts were cut and glued into 24-well tissue culture plates and then sterilized using ethanol. The microwells were then coated with 5% Pluronic Acid for at least an hour and washed with PBS before cell seeding. A single cell suspension of aggregation media containing base media and T0 cytokines supplemented with ROCK inhibitor Y-27632 was then seeded into the wells and allowed to aggregate overnight after centrifuging at 200 × g. Cells are maintained in hypoxia at 37°C in a 5.0% CO_2_ and 5.0% O_2_ atmosphere. After 24 hours, hEB were formed and aggregation media was exchanged for T1 media. On day 4, hEB were removed from AggreWells and placed in Low cluster 6-well plates (NUNC). Corresponding media for T4, T8, T12 was freshly made and exchanged. On T12, cells were returned to normoxia at 37°C in a 5.0% CO_2_ atmosphere. Media was replaced every 8 days onward.

For select experiments, cardiomyocytes were differentiated from human induced pluripotent stem cells (iPSC; ThermoFisher) using modifications to the monolayer-based differentiation protocol carried out by Lian et al. and licensed from the Wisconsin Alumni Research Foundation ^31^. Briefly, iPSCs were seeded on PLO-laminin (Sigma; ThermoFisher) coated tissue culture plates at ∼40,000 cells/cm^2^ in mTeSR1 (StemCell Technologies) until confluency reached 85-95%. Cells were then treated with 12 µM CHIR 99021 (Tocris) in RPMI-B27 basal medium (ThermoFisher) for 22-26 hours, before CHIR was replaced by RPMI/B27 basal medium. After 48 hours in basal medium, cells were treated with 5 µM IWP2 (Tocris) and a potent and proprietary selective tankyrase inhibitor (IC_50_ <10 nM in DLD-1 epithelial cells) for 48 hours in RPMI/B27 basal medium. IWP2 and tankyrase inhibitor were then removed and cells were grown in RPMI/B27 basal medium with insulin with medium change every other day until harvest. Cells were harvested on day 11 to day 15 post-differentiation using TrypLE (ThermoFisher). The percentage of cTnT-positive cells were determined using flow cytometry as reported previously ^31^.

iPSC-differentiated cardiomyocytes (iPSC-CM) were maintained in RPMI-1640 with GlutaMAX and HEPES (ThermoFisher) supplemented with B27 containing insulin (ThermoFisher) and 1x antibiotic/antimycotic (ThermoFisher). Post-differentiation, iPSC-CM were cryopreserved at 2-4 E7 cells/mL in growth media supplemented with 20% dimethylsulfoxide (Sigma).

iPSC-CM were thawed 7 days prior to CaMiRi seeding. T75 flasks (Nalgene) were coated with 25 µg/mL Synthemax II-SC (Corning) overnight at 37°C. Synthemax solution was then aspirated immediately prior to cell seeding. Cryopreserved iPSC-CM were thawed in at 37°C water bath for 4 minutes then transferred to a 50 mL conical tube containing 5 mL of growth media. Cells were centrifuged at 220 × g for 3 minutes and cell pellet was resuspended in 10 mL growth media per T75 flask. After 48 hours, a half media exchange was performed; thereafter, media was fully replaced every 48-72 hours.

### Human cardiac fibroblasts

Human cardiac fibroblasts (hCF) were purchased commercially (PromoCell) and maintained in Advanced DMEM (Life Technologies) supplemented with 10% fetal bovine serum (Sigma), 100 µg/mL basic fibroblast growth factor, or bFGF (Corning), 1x L-glutamine (Gibco), and 1x antibiotic/antimycotic (ThermoFisher). Cells were thawed and maintained according to vendor-provided specifications. bFGF was spiked into media at every media change (2-3 days) to maintain growth factor stability and efficacy. Fibroblasts were cultured in T175 flasks (Nalgene) for up to 10 passages before being discarded.

### Seeding and cultivation of CaMiRi

hPSC-CM were suspended in a collagen mastermix and seeded into cardiac microtissue wells at desired density. Microwell substrates were prepared by sterilizing with ethanol, washing and coating with 5% Pluronic Acid for at least an hour each. During coating, hESC- or iPSC-CM and hCF were prepared. Aggregates from hESC-CM differentiation were incubated in Collagenase for 3 hours at 37°C, 5% CO_2_. Aggregates were then immersed in wash solution (10% FBS and 90% DMEM/F12) and triturated ∼10 times. Once aggregates were dissociated into single cells, samples were counted. iPSC-derived CM and hCF were dissociated from T75 flasks by first washing with phosphate-buffered saline without calcium nor magnesium (PBS^-/-^) followed by a brief rinse with TrypLE (ThermoFisher); cells were then incubated for 10-20 minutes at 37°C until dissociated and counted. The collagen mastermix was prepared by combining the following: 10X M199 (GIBCO), Glutamax (GIBCO), Collagen 1 (3.66 mg/mL) (BD), Glucose (0.3 g/mL) (GIBCO), NaOH (SIGMA), NaHCO3 (0.075 g/mL) (SIGMA), Hepes (GIBCO), GFR Matrigel (BD), ddH20 at appropriate ratios for desired collagen concentrations. Collagen mastermix was maintained on ice under 4°C to prevent premature crosslinking. Finally, 12 uL of mastermix was pipetted into each well (of 96-well plate), after which the entire plate was placed into a normoxic incubator for 15 minutes. After 15 minutes, 300 µL of cell culture media was slowly added to each well so as to not disrupt the polymerized collagen microtissues. Media was exchanged every 2 (iPSC-CM) or 4 (hESC-CM) days. CaMiRi remodeled between 1-3 days depending on input cell composition. Imaging of CaMiRi was performed *in-situ*. Samples were fixed, permeablized, and stained inside the microwells and imaged using a fluorescence microscope.

### Automated Seeding of CaMiRi

A Bravo automated liquid handling platform (Agilent) equipped with a 9-pod deck was utilized for automated seeding of CaMiRi. 2 pods were outfitted with Thermoshake (Inheco) units maintained at 4°C in order to keep source and destination plates cooled to prevent premature gelation of mastermix. Mastermix, containing cells, was evenly distributed to wells E9:H12 of a 96 well U-bottom plate (Falcon). A custom protocol was then employed to distribute 4× 2.5 µL drops of mastermix from the source plate to the north, east, south, and west positions of 16 destination wells at a time; this was repeated 6 times to seed the entire plate and helped minimize dead volume. The destination plate was then centrifuged at 400 × g for 5 minutes at 4°C to drive the mastermix down into a continuous ring at the bottom of each well. Plate was then incubated for 30 minutes 37°C, after which 300 µL of maintenance media, comprised of DMEM/F12 (Life Technologies), 2% FBS (Sigma), 10 µg/mL insulin (Life Technologies), and 1x antibiotic/antimycotic (ThermoFisher), was slowly added to all wells. Plate was returned to 37°C, 5% CO_2_ incubator.

### Video Acquisition and analysis

Throughout the 14 day culture period, images of CaMiRi were acquired every 2 to 3 days using a 4× objective on an Olympus CKX41 microscope equipped with an OptixCam Summit OCS-3.0 digital microscope camera. On day 14, 20 s long videos were acquired using 4× and 10× objectives for each CaMiRi well. Cantilever deflection and contraction frequency were determined by direct measurement or segregating the 10× videos into images and importing them into a custom-designed cantilever tracking software developed in collaboration with CellScale (Waterloo, Canada).

### Automated video acquisition and analysis

Automated videos of CaMiRi-induced post deflections in 96 well plates were acquired at 20x on an Image Xpress Micro 4 (Molecular Devices) by employing a custom journaling solution in Metamorph (Molecular Devices). CaMiRi were maintained at 37°C and 5% CO_2_ during experiments. Briefly, wells were initially imaged at 2x and image analysis was used to identify the x- and y-coordinates of the left post in each well (previous experiments supported the need to only image a single post due to like measurements from both posts; data not shown); magnification was then increased to 20× and a contrast-maximizing autofocus routine was used to focus on the top edge of each post. 20 s videos were then acquired at 100 frames per second using a limited region of interest that captured post displacement. For experiments involving compound addition, compounds were prepared at 5x in a separate U-bottom 96 well plate (Falcon) and maintained on the system’s incubated stage at 37°C. Following acquisition of baseline post deflection, 50 µL of compound was added from source well (one well at a time) to destination well containing 200 µL of medium. Compounds were incubated with CaMiRi for 2 minutes prior to acquisition of an additional 20 s, post-compound addition video.

To extract contractility measurements, videos were converted to 2-dimensional kymographs within Metamorph by projecting each video’s center row of pixels over time in the y-direction. Kymographs were then analysed with a custom MATLAB (Mathworks) script to extract beats per minute, contraction amplitude, contraction velocity, and relaxation velocity.

### Immunohistochemistry and confocal imaging

CaMiRi were rinsed with phosphate buffered saline (PBS) and fixed with 2% paraformaldehyde (PFA) overnight at 4°C. They were then rinsed again with PBS and permeabilized in 0.1% Triton-X in blocking solution (2% BSA in PBS) for 5 min. Microtissues were then incubated in blocking solution for 30 min. Primary antibody was diluted in a 1% BSA in PBS solution and was added and left for 3 days at 4°C. Mouse IgG1 anti-cardiac troponin (Thermofisher MS-295-P) was used at a dilution of 1:100, while Rabbit IgG anti-Vimentin (Abcam AB16700) was used at a dilution of 1:500. After three consecutive washes in PBS of 1 hour each, secondary antibodies were applied for 1 day at 4°C. Alexa Fluor® 488 goat anti-mouse IgG1 and Alexa Fluor® 555 goat anti-rabbit (Life Technologies) antibodies were used at a dilution of 1:1000 for cardiac troponin and vimentin respectively. After the removal of the secondary antibodies, nuclear staining was performed using a DAPI solution (1 µg/mL in PBS) for 15 min. The microtissues were then washed in PBS (3 × 1 hour) before mounting on glass slides using fluoro-safe mounting media (Dako, S3023). Mounted samples were allowed to cure for 1 day and then imaged using a confocal microscope (Zeiss LSM700 Confocal Microscope).

### Statistical analysis and data representation

Statistical analysis for the Central Composite Design (CCD) experiment was performed with Design Expert v.9 software (Stat-Ease Inc.). The CCD was used to determine the optimal tissue formulation for CaMiRi based upon 3 parameters of collagen concentration, total cells per tissue, and percentage of cardiomyocytes and cFB in the tissue. The two factors used as metrics were contraction frequency and contractile force (as measured by cantilever deflection). The design consisted of 64 experimental runs with four replicates at the factorial and axial points, and eight replicates at the centre point (Figure 3A). The experimental design and data analysis were performed using Design Expert v.9 (Stat-Ease Inc.). Statistical analyses between groups were computed by an ANOVA using PSPP v.0.8.3 software, with pairwise comparisons performed by Tukey’s post-hoc test. A quadratic model was selected to fit the data and an ANOVA was performed to assess the significance of each term as a means of maximizing successful wire formation and cantilever deflection. Statistical analysis for the drug screen experiment was conducted with pairwise comparisons performed by the Mann–Whitney U test. All error bars represent the standard deviation of three or more biological replicates. Asterisks (*) indicate statistical significance between conditions of *P* < 0.05. All data analyses, including graphical representations, were performed using Excel (Microsoft, Redmond, WA).

---

**Supplementary Figure 1.**

Dimensions of well and microcantilever design within 96-well plate. (A) Dimensions of microcantilever geometries as shown in cross-sectional view. (B) Higher detail dimensions of individual well. (C) Total volume and surface area calculations for all compartments of well. (D) Beam bending formula for calculating force versus displacement curve. Deflection (δ), Force (F), Length of cantilever (L), Young’s Modulus (E), Moment of inertia (I), Radius of cylinder (r).

**Supplementary Figure 2.**

Drafting of 3D printed mold shows decreased areas of stress during demolding process. (A) Green areas represent areas of high stress during demolding process, in contrast to yellow walls which represent low stress during demolding. Drafting angles into the walls convert green high stress areas into yellow low stress areas (left vs. right panels). (B) Schematics of angled walls show degree of drafting integrated into CAD design before 3D printing.

**Supplementary Figure 3.**

Cell seeding of CaMiRi. (A) Seeding and tissue formation process flow. 12μl of cell-laden matrix are pipetted into the reservoir of each well, as shown in the first diagram. Polymerize in the incubator for 20 minutes. 200 uL of cell culture media is then slowly added into each well over top of the polymerized cell-laden collagen gel, as shown in the second diagram and return to the incubator. Over time the cells remodel around the two cantilevers as the cells start to form a compact tissue. (B) Time lapse imaging of remodeling CaMiRi composed of NKX2-5-GFP+ hPSC-CM. (C) Top and side view of cardiac microtissue on microcantilevers. CaMiRi in bright field shown on left, and in red (due to cell tracker dye) shown on right. Scale bars represent 2 mm.

**Supplementary Figure 4.**

Step-by step visual process of plate fabrication and cell seeding. (A) 3D printed positive mold of CaMiRi platform. (B) PDMS negative mold made of previous 3D printed mold. (C) PDMS positive molded from previous step. (D) Polyurethane negative master mold cast using PDMS positive mold from previous step. (E) PDMS CaMiRi plate molded from polyurethane master mold from previous step and autoclaved, ready to be coated and seeded. (F) Cell-laden collagen pipetted into wells over ice. (G) Excess cell-laden collagen on plate walls. (H) After gently tapping plate on flat surface, cell-laden collagen has completely sunk into the reservoirs to create an annulus of cell-laden collagen. (I) After 20 minutes in incubator, CaMiRi have polymerized. 200 mL of media is slowly pipetted into each well using a multichannel pipettor.

**Supplementary Figure 5.**

Video capture and analysis of deflecting cantilevers. (A) Schematic depicting camera located above well capturing video of deflecting cantilever and edge detection (B) to calculate pixel movement during contractions and (C, D) batch processing of data using custom software designed by CellScale.

**Supplementary Figure 6.**

Central Composite Design (CCD)-informed formulation of CaMiRi. (A, B) Legend and schematic of CCD parameters and conditions tested (low, medium, and high values) chosen for percentage of cardiomyocytes and cardiac fibroblasts in tissue (50%, 70%, 90% cardiomyocytes), collagen concentration (1.8, 2.0, and 2.2 mg/mL), and cell number per tissue (40,000; 70,000; 100,000 cells per tissue). (C) Contraction frequency of cardiac microtissues in all conditions measured in beats per minute show no significant differences between conditions. (D) Summary of surface functions of each combination of 3 factors tested in CCD study result in a predicted best-case formulation of 90% CM, 2.0 mg/mL collagen 1 concentration, and 75,000 cells per CaMiRi.

**Supplementary Figure 7.**

Central Composite Design (CCD)-informed formulation of CaMiRi. (A) Schematic of CCD parameters (low, medium, and high values) chosen for collagen concentration (1.8, 2.0, and 2.2 mg/mL), cell number per tissue (40,000; 70,000; 100,000 cells per tissue), and percentage of cardiomyocytes and cardiac fibroblasts in tissue (50%, 70%, 90% cardiomyocytes). (B) Surface functions of each combination of 3 factors tested in CCD study resulting in a predicted best case formulation of 90% CM, 2.0 mg/mL collagen 1 concentration, and 75,000 cells per CaMiRi.

**Supplementary Figure 8.**

Electrophysiology-based assay using CaMiRi. (A, B) Electrophysiological assessment of maximum capture rate (a measure of tissue response to stimulation), and excitation threshold (a measure of tissue excitability), of growth factor effects on cardiac microtissues over 2 weeks. IGF-1 and HRG show improved excitation threshold in cardiac microtissues. Data are reported from three experiments with at least two replicates each, as the mean ± standard deviation. *P < 0.05.

**Supplementary Figure 9.**

Automated seeding of CaMiRi. (A) 4× 2.5µL drops of cellular mixture were pipetted to N, E, S, and W positions inside each well. (B) 16 wells were seeded at a time, in 6 iterations to cover the 96 well plate, using an Agilent Bravo automated liquid handler. (C) Discrete droplets post-dispense (left) were driven into contiguous rings (right) following cooled centrifugation for 5 minutes at 200 × g. (D) Centrifugation conditions were optimized functionally by measuring CaMiRi contraction amplitude 14 days post-seeding. (E) 96 well plate of CaMiRi, 14 days after automated seeding.

**Supplementary Figure 10.**

Layered multicellular CaMiRi. (A) Using a layering approach via sequential seeding, multicellular tissues can be generated using the CaMiRi platform. Cell type 1 (green) can be seeded first, permitted to remodel, and then cell type 2 (red) can be seeded subsequently to create an additional outer layer.

