## Supplementary Figures for "Functional arrays of human pluripotent stem cell-derived cardiac microtissues"

Supplementary Figure 1

A

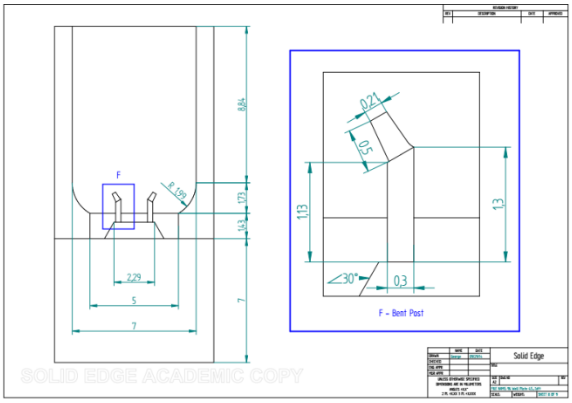


B

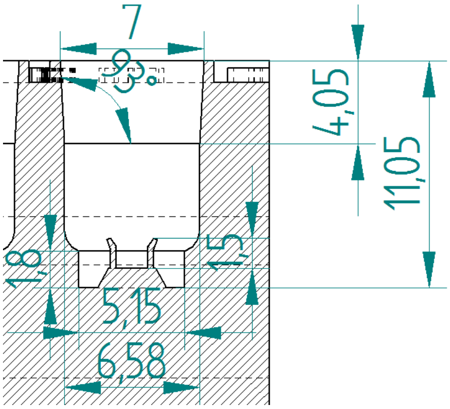


C

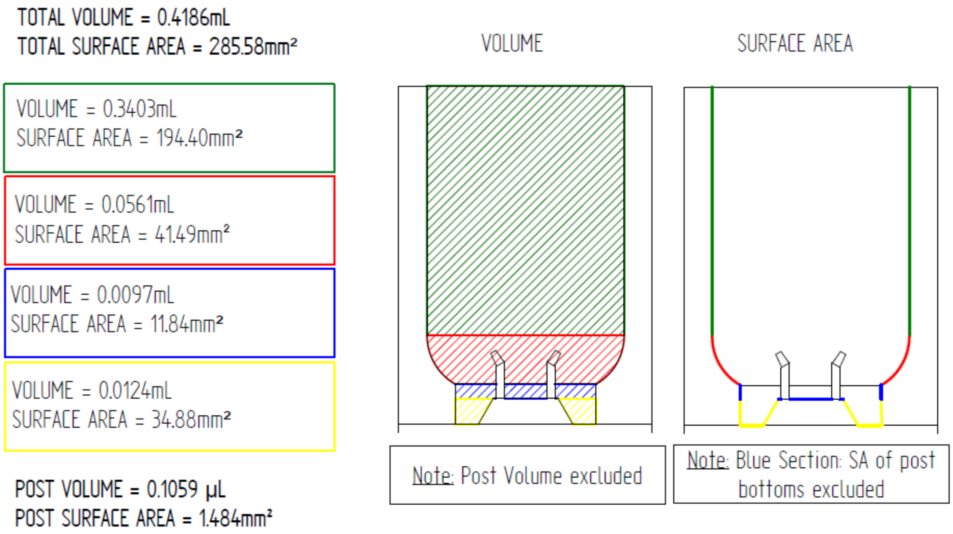


D

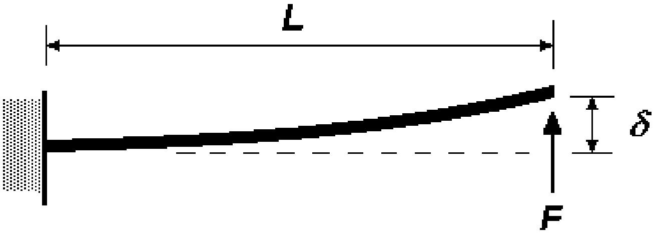


$$\delta = FL^3 / 3EI$$

$$I = \pi r^4 / 4$$

### Supplementary Figure 2

**A**

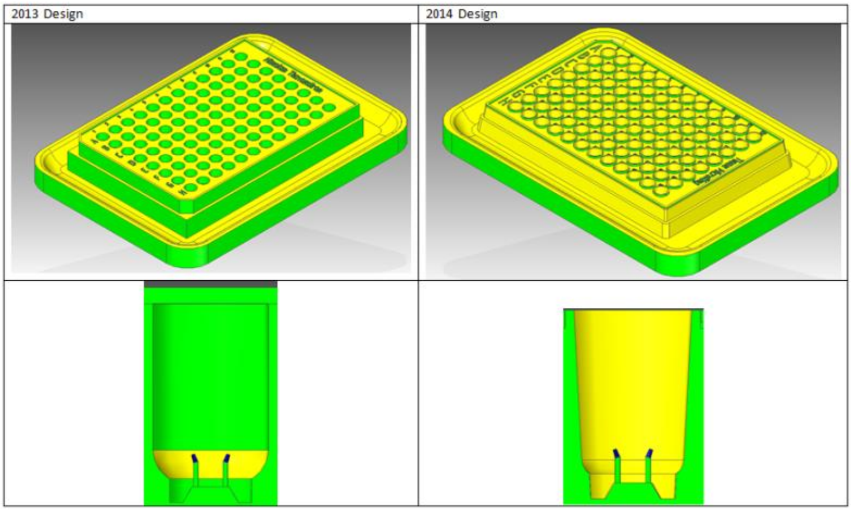


**B**

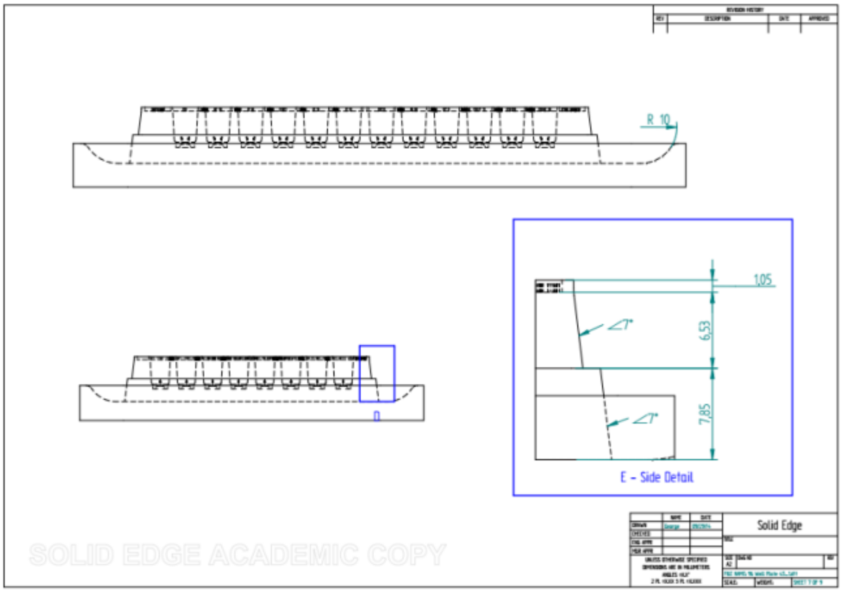

### Supplementary Figure 3

**A**

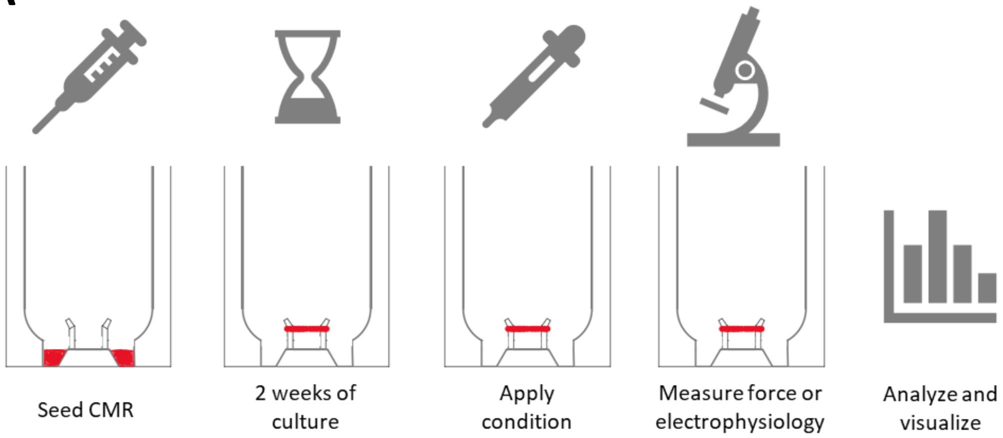


**B**

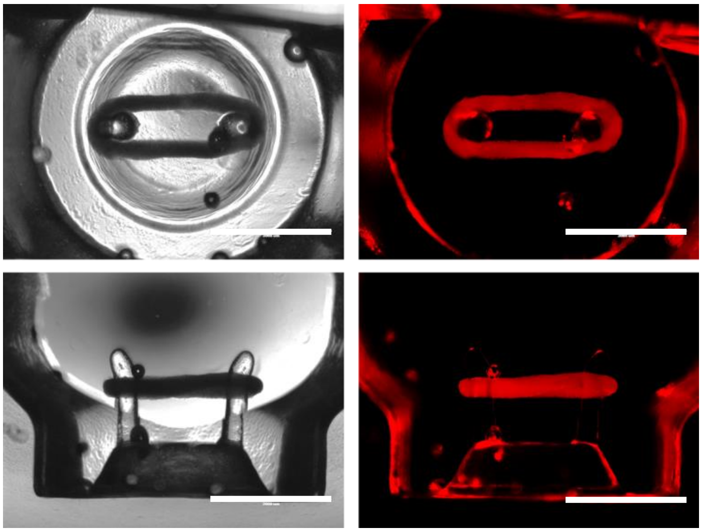

Supplementary Figure 4

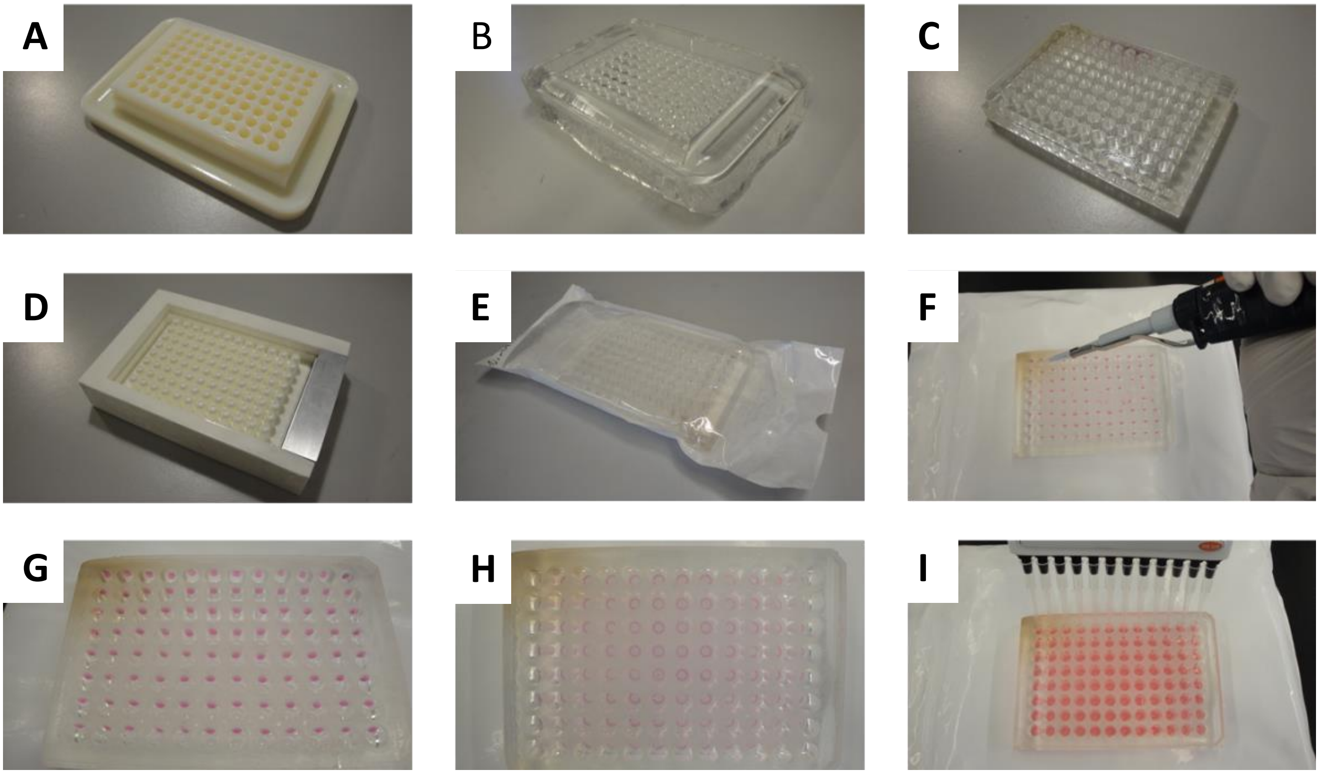

### Supplementary Figure 5

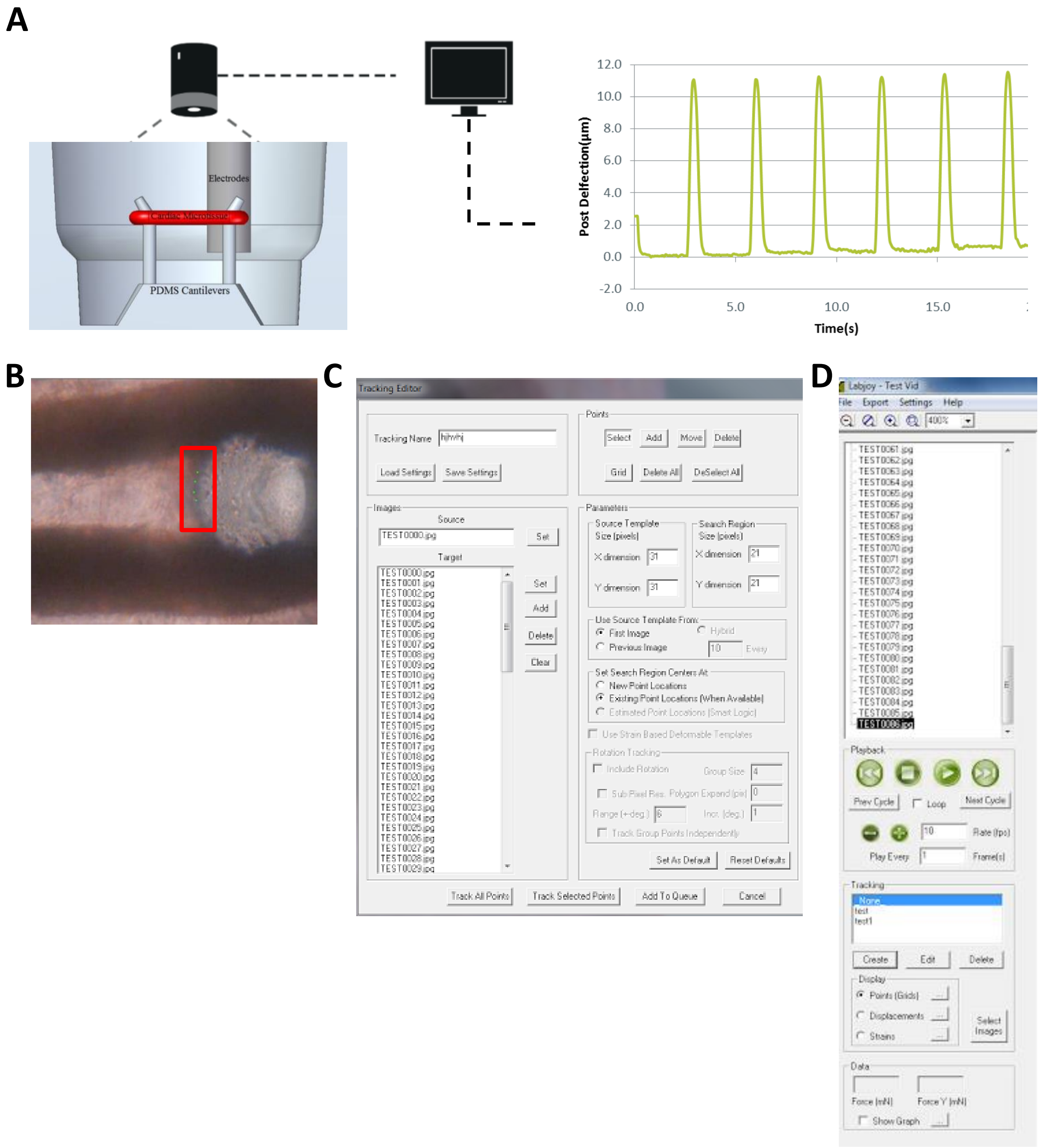

### Supplementary Figure 6

**A**

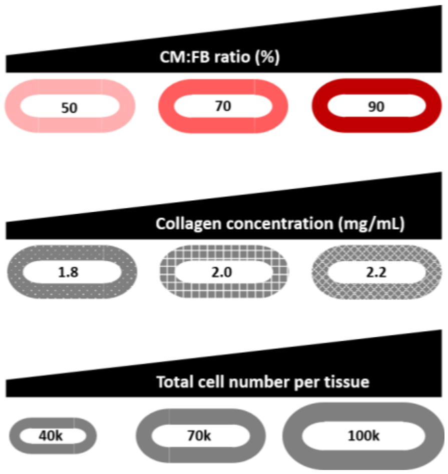


**B**

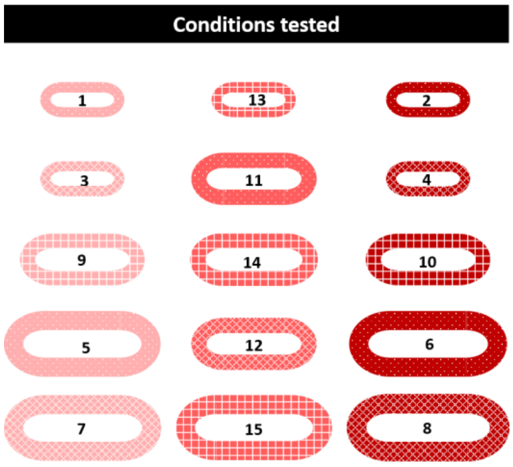


| Condition | % CMs | Collagen | Cell Number |
|---|---|---|---|
| **1** | 50 | 1.8 | 40,000 |
| **2** | 90 | 1.8 | 40,000 |
| **3** | 50 | 2.2 | 40,000 |
| **4** | 90 | 2.2 | 40,000 |
| **5** | 50 | 1.8 | 100,000 |
| **6** | 90 | 1.8 | 100,000 |
| **7** | 50 | 2.2 | 100,000 |
| **8** | 90 | 2.2 | 100,000 |
| **9** | 50 | 2.0 | 70,000 |
| **10** | 90 | 2.0 | 70,000 |
| **11** | 70 | 1.8 | 70,000 |
| **12** | 70 | 2.2 | 70,000 |
| **13** | 70 | 2.0 | 40,000 |
| **14** | 70 | 2.0 | 100,000 |
| **15** | 70 | 2.0 | 70,000 |

**C**

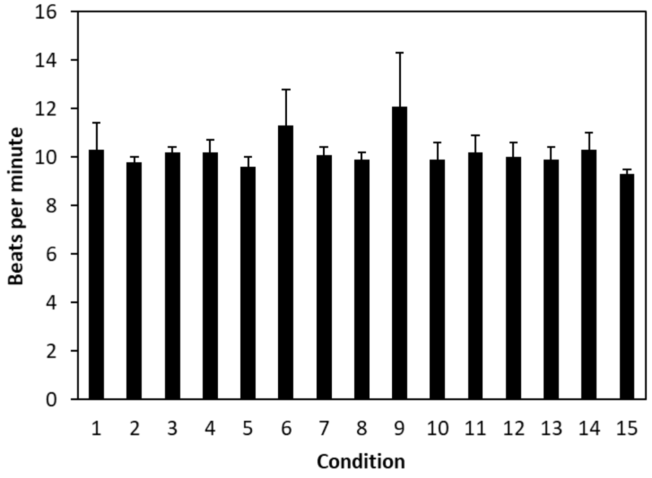


**D**

| **Predicted best case:** | |
|---|---|
| Seeding cell ratio | 90% CMs |
| Collagen concentration | 2 mg/mL |
| Cell number per well | 75,000 cells |

Supplementary Figure 7

**A**

| | % CMs | Collagen | Cell Number |
|---|---|---|---|
| 1 | 50 | 1.8 | 40,000 |
| 2 | 90 | 1.8 | 40,000 |
| 3 | 50 | 2.2 | 40,000 |
| 4 | 90 | 2.2 | 40,000 |
| 5 | 50 | 1.8 | 100,000 |
| 6 | 90 | 1.8 | 100,000 |
| 7 | 50 | 2.2 | 100,000 |
| 8 | 90 | 2.2 | 100,000 |
| 9 | 50 | 2.0 | 70,000 |
| 10 | 90 | 2.0 | 70,000 |
| 11 | 70 | 1.8 | 70,000 |
| 12 | 70 | 2.2 | 70,000 |
| 13 | 70 | 2.0 | 40,000 |
| 14 | 70 | 2.0 | 100,000 |
| 15 | 70 | 2.0 | 70,000 |

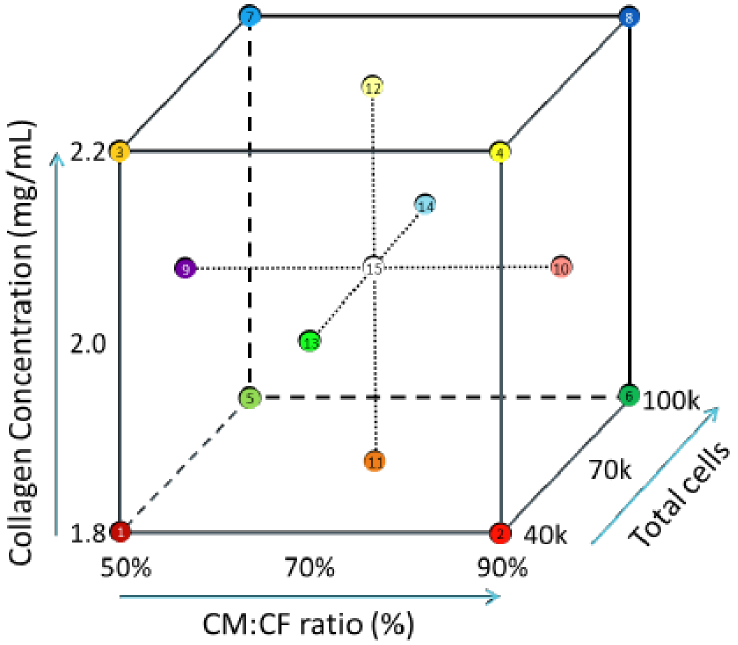


**B**

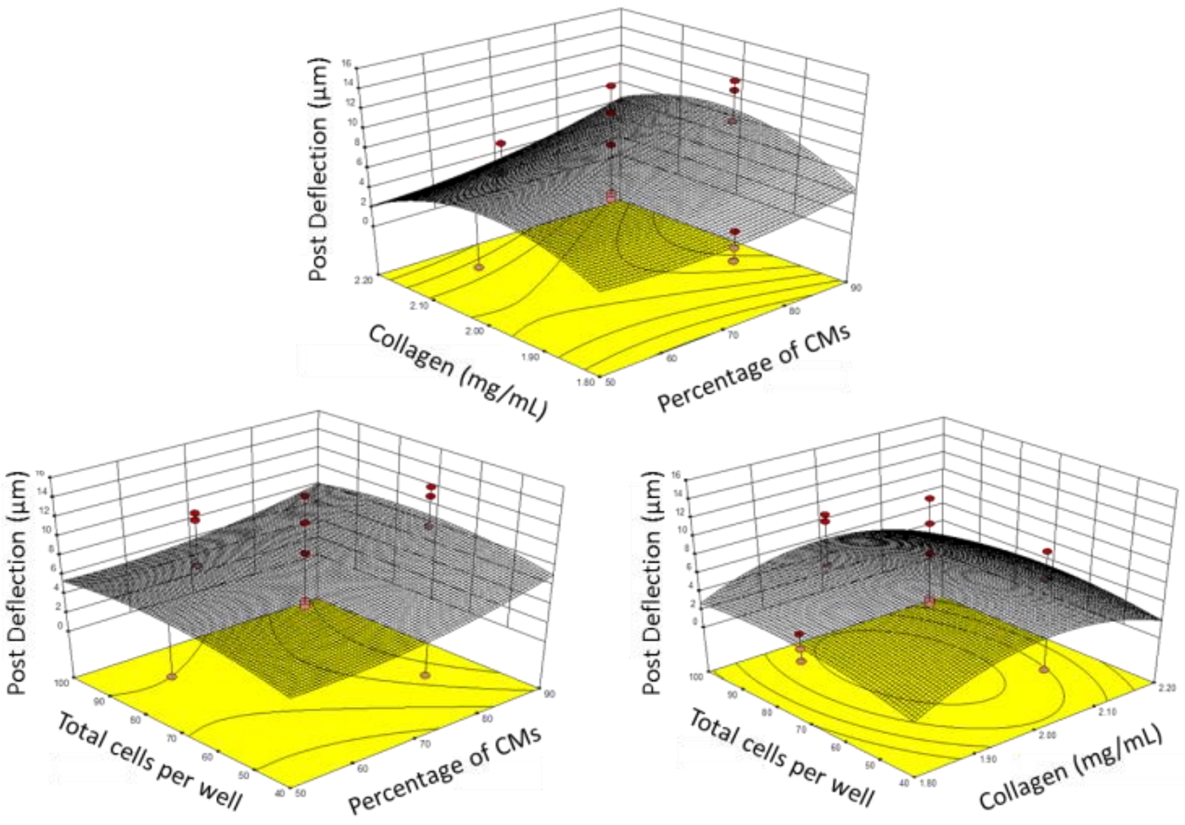

Supplementary Figure 8

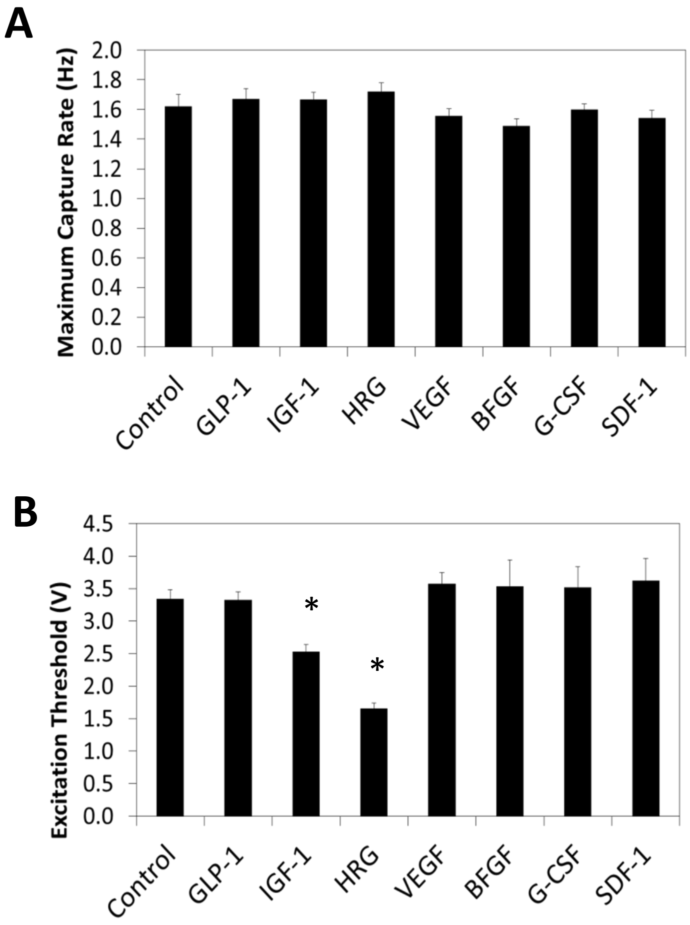

### Supplementary Figure 9

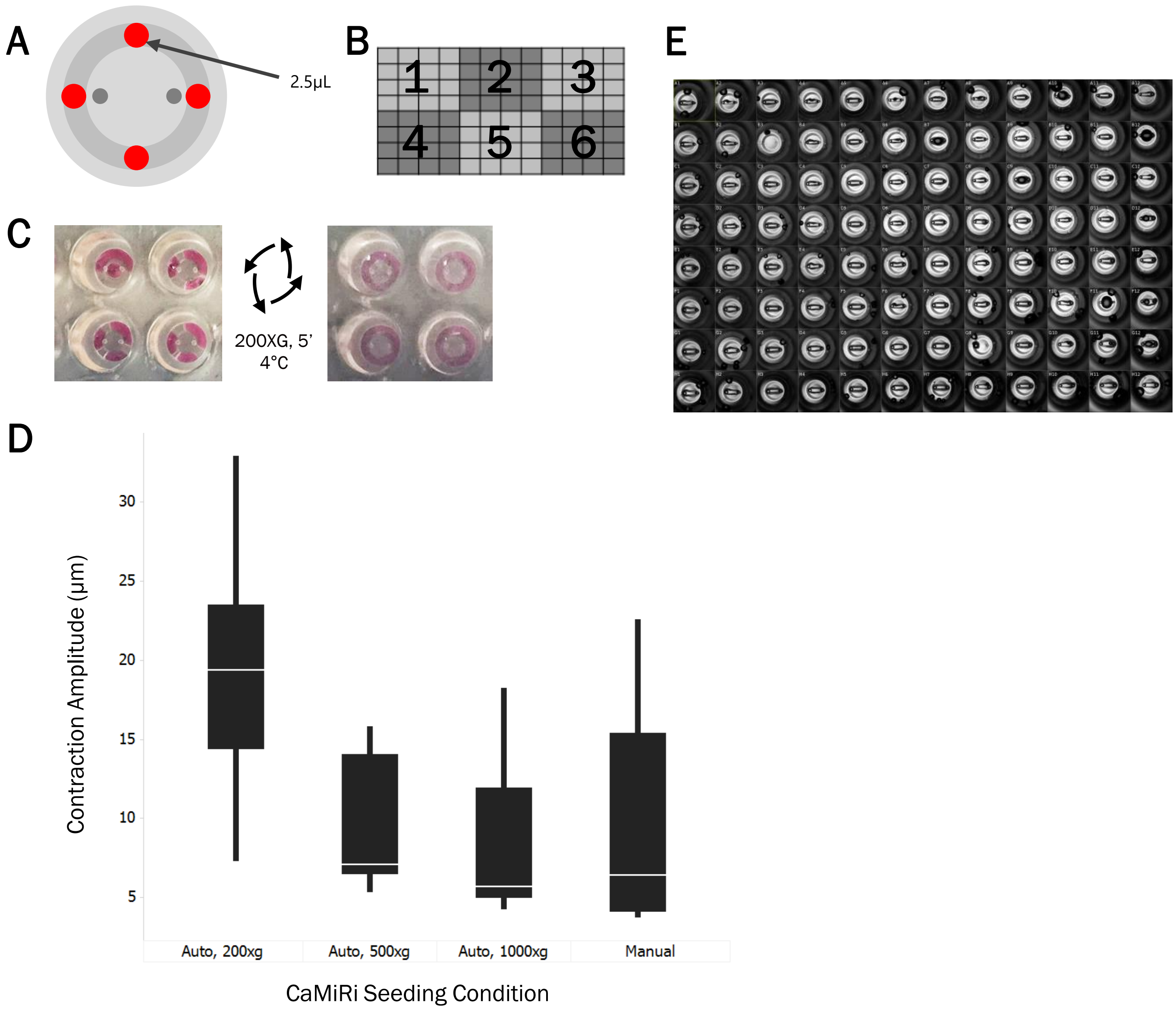

Supplementary Figure 10

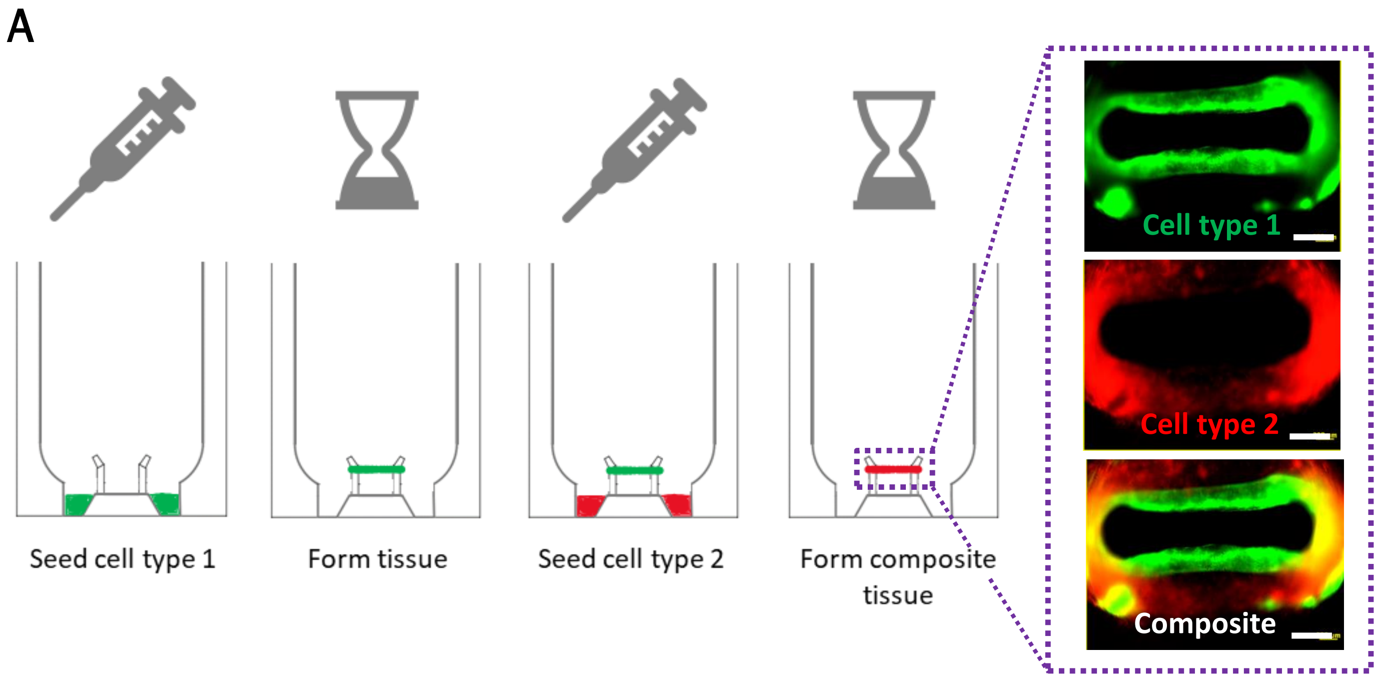
